# Quasi-Periodic Patterns of Intrinsic Brain Activity in Individuals and their Relationship to Global Signal

**DOI:** 10.1101/180596

**Authors:** Behnaz Yousefi, Jaemin Shin, Eric H. Schumacher, Shella D. Keilholz

## Abstract

Quasiperiodic patterns (QPPs) as reported by Majeed et al., 2011 are prominent features of the brain’s intrinsic activity that involve important large-scale networks (default mode, DMN; task positive, TPN) and are likely to be major contributors to widely used measures of functional connectivity. We examined the variability of these patterns in 470 individuals from the Human Connectome Project resting state functional MRI dataset. The QPPs from individuals can be coarsely categorized into two types: one where strong anti-correlation between the DMN and TPN is present, and another where most areas are strongly correlated. QPP type could be predicted by an individual’s global signal, with lower global signal corresponding to QPPs with strong anti-correlation. After regression of global signal, all QPPs showed strong anti-correlation between DMN and TPN. QPP occurrence and type was similar between a subgroup of individuals with extremely low motion (or even high motion) and the rest of the sample, which shows that motion is not a major contributor to the QPPs. After regression of estimates of slow respiratory and cardiac induced signal fluctuations, more QPPs showed strong anti-correlation between DMN and TPN, an indication that while physiological noise influences the QPP type, it is not the primary source of the QPP itself. QPPs were more similar for the same subjects scanned on different days than for different subjects. These results provide the first assessment of the variability in individual QPPs and their relationship to physiological parameters.

## Introduction

Spontaneous fluctuations in the blood oxygenation level dependent (BOLD) MRI signal attributed to intrinsic brain activity exhibit varied dynamic patterns that have been described as time-varying correlation (Chang and Glover 2010; Allen et al., 2012; Keilholz et al., 2013), sparse patterns of localized spontaneous activations (Petridou et al., 2013; Caballero Gaudes et al., 2013), or co-occurring activity (Liu and Duyn 2013; Chen et al., 2015), and repeated spatiotemporal patterns (Majeed et al., 2011; Kiviniemi et al. 2016; also see Preti et al., 2016 for a recent review on varied dynamic patterns). Quasi-periodic patterns (QPPs) fall into the last category and consist of a reproducible pattern of spatial changes that repeat over time, exhibiting alternation of high and low activity in particular areas and propagation of activity along the cortex. QPPs were first observed in anesthetized rats as a bilateral propagation of high activity from lateral to medial cortical areas, followed by an echoing propagation of low activity (Majeed et al., 2009). Majeed and colleagues (2011) subsequently developed a pattern-finding algorithm to identify QPPs in humans, which involved alternating high and low activity in default mode (DMN) and task positive (TPN) networks. Animal and human studies have shown that QPPs are linked to infra-slow (<0.1Hz) electrical activity (Pan et al., 2013; Thompson et al., 2014a, 2014b, 2015; Keilholz 2014; Grooms et al., 2017) and represent a different type of activity than the higher frequency activity tied to time-varying BOLD correlation between areas (Thompson et al., 2015; Keilholz et al., 2016). Although the infra-slow electrical signals themselves are still poorly understood, possibly arising from coordinated interactions between neurons, glia, and the vasculature (Keilholz et al., 2016; Thompson et al., 2014a), nevertheless, infra-slow activity is one of the best candidates for the coordinating mechanisms within and between brain’s large-scale networks (see discussion in Thompson et al., 2014a, 2014b for literature review). Hence BOLD QPPs may also reflect aspects of such mechanisms. Like traditional BOLD-based networks of functional connectivity, QPPs have been observed in mice (Belloy et al., 2017), rats (Pan et al., 2013; Thompson et al., 2014a, 2014b; Magnuson et al., 2010), monkeys (Abbas et al., 2016a), and humans (Majeed et al., 2011; Kiviniemi et al., 2016), in states ranging from deeply anesthetized to awake, again suggesting that they represent a fundamental aspect of the brain’s functional organization.

The QPP algorithm is a correlation-based iterative algorithm that identifies a recurring spatiotemporal template during a functional MRI scan (Majeed et al., 2011). First, a segment of a preset number of consecutive timepoints is selected based on a random starting point. Sliding correlation of this segment with the functional scan is calculated, timepoints corresponding to local maxima above a preset threshold are selected, and segments of the scan starting at those timepoints are averaged together with the original segment to create a template. The sliding correlation is then repeated with the template in place of the initial random segment and the process repeats until the template exhibits negligible change between iterations **(Fig. S1).** Thus far, the QPP method has been used with multiple randomly selected starting timepoints followed by a hierarchical clustering to select the most representative QPP. It has typically been applied to concatenated scans from all subjects, meaning that a single template is derived for the entire group, with variable levels of contribution across subjects.

To examine variability in the QPPs at the individual level, we made three modifications to the QPP method that increased its robustness: 1) segments starting at all timepoints of each scan are used to create templates rather than using a limited number of random timepoints, 2) a customized criterion for the selection of the most representative QPP is introduced based on maximizing the template’s correlation with functional scan and the template’s periodicity, 3) a method for phase-adjusting a QPP is introduced in order to correctly compare QPPs of the same subject in different days or QPPs of different subjects. The modified algorithm was applied to resting state functional MRI data from the Human Connectome Project (HCP) (Van Essen et al., 2013; Glasser et al., 2016a) at the individual level. To examine the effects of motion, analysis was performed on a subgroup of 40 individuals with the lowest motion and compared to 470 subjects with more moderate levels.

Large-scale patterns such as QPPs are likely to contribute to the global signal, so analysis was performed before and after global signal regression. Slow variations in respiration depth and rate and heart rate have been shown to correlate with variations in the global signal (Power et al., 2017; Liu et al., 2017; Keilholz et al., 2016) and with variations in the default mode network (Birn et al., 2006 and 2008; Chang and Glover 2009; Change et al., 2009), so the relationship between respiration and heart rate variation and QPPs was also examined. To compare the levels of variability within individuals to variability across individuals, the HCP resting state scans acquired over two subsequent days were analyzed separately. This report provides the first examination of individual variability in QPPs and provides further support that the patterns reflect neural activity rather than physiological noise or motion. Because QPPs contribute substantially to functional connectivity, especially in the default mode network, a better understanding of their properties and sources can provide insight into the connectivity differences underlying different behavioral states and traits and connectivity changes associated with neurological and psychiatric disorders.

## Method

### Doto and preprocessing

Minimally preprocessed grayordinate and FIX de-noised resting state scans were downloaded from the Human Connectome Project S900 release (Glasser et al., 2013; Glasser et al., 2016a). To minimize potential contributions from motion, the head motion data from the 820 HCP subjects with four complete resting state scans was inspected and 40 subjects with mean frame-wise displacement (FD; Power et al., 2014) less than 0.12mm per scan for all four scans were selected for initial analysis, designated "low movers" in the text. After QPPs proved detectable and robust at the individual level in this group of high quality data, the selection criterion was relaxed to include 470 subjects with temporal ratio of FD>0.2mm less than 0.4, or equivalently FD-spikes < %40, per scan for all the four scans, designated the "low-moderate movers" in the text (mean FD < 0.2mm; see **Fig. S2a).** Note that all low movers were included among the low-moderate movers. Previous work that has specifically examined motion in the HCP dataset used spikes in temporal Derivative then rms VARiance over elements (DVARS or DV) in addition to FD to identify high motion timepoints (Burgess et al., 2016; Siegel et al. 2016). Therefore, we examined DV as well **(Fig. S2a** and **S2b).** In the initial analysis of the 40 low movers, QPP detection was performed using the timeseries of all ~92K cortical vertices and subcortical voxels. To maintain practical computation times for the larger group of low-moderate movers with our modified approach to QPP detection, the spatial dimension was reduced to 360 cortical parcels (Glasser et al., 2016b) by averaging vertices’ timeseries across each parcel and normalizing the z-scores of each parcel’s timeseries.

The timeseries of each vertex/voxel per scan underwent the following processing steps: 1) demeaning 2) band pass filtering (Butterworth, fourth order, 1dB cutoff frequencies: 0.01 and 0.1Hz, using Matlab fdesign and filtfilt functions, 424 zeros pads were inserted at both ends of each 1200-timepoint scans before filtering and were removed afterwards to minimize transient effects), and 3) normalization by their own standard deviations. For the initial analysis, no global signal regression was performed.

### Modified QPP detection method

The original approach described by Majeed et al., to identify QPPs involved choosing a limited number of randomly selected starting timepoints, running the main algorithm and finding the templates corresponding to those starting timepoints, using hierarchical clustering on those templates, and finally selecting the template that has the maximum average correlation with the rest of the templates in the biggest cluster. However, some starting timepoints result in a template that has a low correlation with the functional scan or does not occur often. Other starting timepoints may pick up the same template but at different phases. To examine the full potential of the main algorithm, a computationally efficient Matlab script was developed to inspect the templates resulting from all the timepoints. It exactly replicated the main algorithm, keeping all free parameters the same: correlation threshold of 0.1 for the first 3 iterations and 0.2 for the rest, maximum iteration of 20. QPPs in humans are approximately 20s long (Majeed et al., 2011); for this study, the window length, or template duration, was set to 30 timepoints (21.6s). Four scans were concatenated for each subject and the new QPP script inspected every timepoint, giving 4x(1200-30+1)=4684 time points.

**Fig. 1** demonstrates the use of all timepoints to identify all possible templates: in **Fig. 1a,** the time course of sliding correlation for a template corresponding to a single starting timepoint is shown for one individual. In **Fig. 1b,** timepoints corresponding to supra-threshold local maxima of such correlation are plotted in the vertical axis for each starting time point, and the value of correlation is indicated by color. When all timepoints are considered as starting timepoints, considerable redundancy results. Sequential starting timepoints or those ~30 timepoints apart are essentially the same template. Also, templates from some timepoints exhibit low correlation or periodicity. Hence, for the template resulting from each timepoint, values of its sliding correlation at local maxima that were above the threshold of 0.2 at the final iteration were summed and the template with the highest sum was designated as the most representative QPP. Selected in this way, the most representative QPP is guaranteed to have high correlation and large numbers of occurrences relative to other templates.

**Figure 1.**
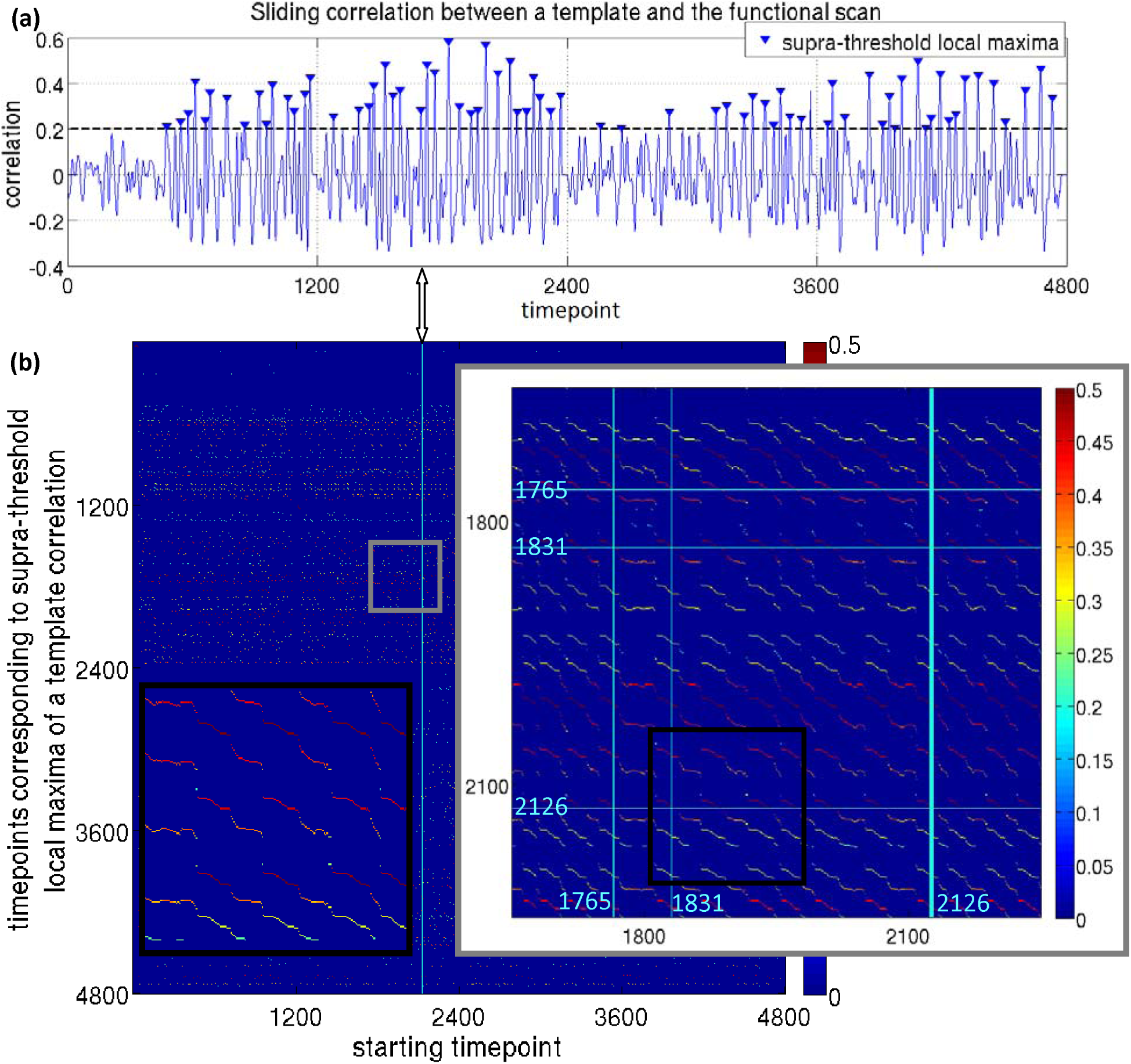
**(a)** Correlation between a template corresponding to a single starting timepoint and the entire scan for one individual, **(b)** Timepoints corresponding to supra-threshold local maxima of a template correlation timecourse (e.g., marked in Fig1a) plotted in the vertical axis for each starting time point, with the value of correlation is indicated by color. When all timepoints are considered as starting timepoints, considerable redundancy results: (i) sequential starting timepoints detect different phases of the same template (black inset), and (ii) starting timepoints which are ~30 timepoints apart are essentially part of the same template; for instance, Fig. 1a corresponds to the starting timepoint 2126 **(thick** vertical cyan line) and 1765 and 1831 are among supra-threshold maxima (and the corresponding template is the average of the segments of scan starting at those maxima after convergence of the algorithm), when 1765 is the starting timepoint, 1831 and 2126 are among supra-threshold maxima and when 1831 is the starting time point, 1765 and 2126 are among supra-threshold maxima.

### Characterizing the spatial extent of the anti-correlation within the duration of a QPP

One of the most prominent features previously noted in the QPP is that patterns of activity are 180 degrees out of phase, i.e., anti-correlated, for regions of the DMN and TPN. For this analysis, we first reduced the dimensionality of the low-mover’s templates by averaging voxels in each parcel to obtain a 360×30 representation of the templates comparable to the parcel-wise templates of the low-moderate movers. To test for and characterize anti-correlation between DMN and TPN, each of 14 parcels in left posterior cingulate cortex (LPCC) was taken as a seed, one at a time, and the histogram of Pearson correlation values between the 30-timepoint template timeseries of each seed with all 360 parcels was calculated with bin centers −0.9:0.2:0.9 (see **Fig. S3** for a complete description of the procedure). The histogram with maximum number of parcels in the first bin was selected for each subject. In the case of zero count for the first bin for all seeds, the timeseries of all seeds were averaged and again a histogram of Pearson correlation values with 360 parcels was calculated with the same bin centers. Not all parcels in Glasser’s PCC region are strongly task negative (Glasser et al., 2016b), which argued against using the average of all seeds for everyone. The seed that maximized the spatial extent of anti-correlation was different across individuals, in line with Chen et al. 2017.

### Relationship between QPPs and global signal

Because large, spatially coherent patterns such as the QPP may contribute to the global signal, the relationship between QPPs and the global signal was explored as one of the central pillars of the current work. For the low movers, we used the average of the timeseries of cortical vertices and subcortical voxels (cortical vertices and subcortical signal) as an approximation for the global signal, since previous studies have shown the global signal originates primarily in the gray matter (Glasser et al., 2016a; Power et al., 2017). For the 470 low-moderate movers, we used the average of the timeseries of cortical vertices (cortical signal) since their QPP analysis was performed using only cortical parcel-wise timeseries. The total cortical and subcortical signal is highly correlated with the cortical signal (r=0.99; Fig. S4).Throughout this article, we have used the term global signal in general statements in place of cortical and subcortical signal or cortical signal.

The following approaches were taken to explore the relationship between QPPs and the global signal:

1) The values of the first and the last bins of the correlation histograms described above, which reveal the spatial extent of strong negative and strong positive correlation in the QPPs, were plotted against the root mean square (rms) of the global signal. Suggested by a clear visual division in the values of the last bin, QPPs were clustered into two groups using Kmeans, with the values of the last bin being the input. The rms of the global signals were compared between groups using an unpaired t-test.

2) Global signal was regressed and the QPP analysis was repeated. For global signal regression, the timeseries of each vertex underwent only the first two steps of the preprocessing, then global signal was calculated by averaging across all vertices. General linear modeling (GLM) was performed with the global signal as the regressor. The residual timeseries were averaged across each parcel, the parcels’ timeseries were z-scored and the QPP analysis was repeated. The correlation histograms were then calculated again to determine the effect on the spatial extent of strong positive and negative correlation.

3) The QPP template of each subject was convolved with its corresponding correlation time course (QPP⊗C) and the result was cross-correlated with the global signal (see Fig. S5 for procedure). Throughout this work, all cross correlation time lags are allowed to range from −20 to 20 timepoints, and, the maximum of the absolute value of cross-correlation and the corresponding lag are reported. The goal of this procedure was to determine whether QPP occurrence coincided with timepoint-by-timepoint fluctuations of the global signal. The temporal pattern of the QPP is different for each parcel (or voxel), so for each subject, we selected the parcel whose raw timeseries had the highest correlation with the global signal (ROI_GS_). To build a null distribution for cross-correlation values, phase-randomized timeseries were made out of QPP⊗C at ROI_GS_ and the global signal as described in Majeed et al., 2011 for each subject. The phase randomized time series were then cross-correlated, and the Kolmogorov-Smirnov (KS) test was adopted to compare the distributions from the real and phase-randomized data.

### Relationship between QPPs and slow physiological variations

Slow variations (<0.1Hz) in respiration rate and depth and cardiac rate are known to correlate with fluctuations of the global signal (Power et al., 2017; Liu et al., 2017; Keilholz et al., 2016) and with BOLD fluctuations in the DMN (Birn et al., 2006; Chang and Glover 2009; Change et al., 2009). To examine the relation between QPPs and slow respiratory- and cardiac-induced BOLD fluctuations, the respiratory belt and cardiac traces of the 470 low-moderate movers: were despiked using Matlab medfilt1, amplitude clipped to −2.5 to +3.5 standard deviation from mean to remove outliers not fixed by despiking, band pass filtered with a fourth order Butterworth with3dB cutoff frequencies of 0.01 and 1Hz for respiratory traces and 0.6 and 3Hz for cardiac traces, using Matlab fdesign and filtfilt functions (zeros pads were inserted at both ends before filtering and were removed afterwards to minimize transient effects), and amplitude rescaled to 0-100(Fig. S6a). For manual quality control, histograms of the preprocessed traces in time (Kasper et al., 2017) and their frequency spectra were visually inspected (Fig. S6b). The number of subjects with acceptable quality respiration and cardiac data for all four scans were 422 and 326, respectively.

Respiratory Variation (RV), defined as the standard deviation of the respiratory trace in a sliding window of 7.2s, which equals ~two respiratory cycles, centered around each fMRI scan timepoint (Chang et al., 2009), was then calculated for each scan. HCP respiration data often has short lapses causing spikes in RV; therefore, based on the histogram of the standard deviation of RV (std RV), only subjects whose std RV for all scans were within three standard deviations above the median were included (404 subjects; Fig. S6c). Heart rate Variation (HV), defined as the average of time between successive peaks of cardiac trace in a sliding window of 7.2s centered around each fMRI scan timepoint (Chang et al., 2009) was also calculated. For peak detection, the Matlab findpeaks function was used with the minimum peak distance set to a value that corresponded to two-thirds of the largest peak in frequency domain (Fig. S6a showcases the effectiveness of this simple method). Based on the histogram of the standard deviation of HV (std HV), only subjects whose std HV for all scans were within three standard deviations above the median were included (315 subjects; Fig. S6c). 292 subjects had both high quality respiration and cardiac data for further physiological related analyses.

The values of the first and the last bin of the correlation histogram of each QPP, corresponding to the spatial extent of strong negative and positive correlations, were plotted against the average of std RV and std HV across four scans. The average of std RVs and std HVs across four scans was compared between groups with the two QPP types using an unpaired t-test. In addition, physiological noise was regressed and QPP analysis was repeated. In order to regress physiological noise, the Respiratory Response Function (RRF) and Cardiac Response Function (CRF), whose analytical forms are introduced in Birn et al., 2008 and Chang et al.,2009 (see Fig. S7), were convolved with RV (RRF⊗RV) and HV (CRF⊗HV), respectively, to build the estimate of respiratory- and cardiac-induced signal fluctuations. RRF⊗RV and CRF⊗HV were separately cross-correlated with parcel-wise raw timeseries to identify the optimal lag that resulted in the maximum correlation per parcel. GLM was performed, per parcel, with a 2-column regressor built from RRF⊗RV and CRF⊗HV, each shifted by their optimal lag for that parcel. Residual timeseries were z-scored, QPP analysis was repeated, and the correlation histograms were again calculated to determine the effect of physiological noise regression on the spatial extent of strong positive and negative correlation.

### Metrics for QPP correlation and periodicity

To obtain a single value that describes the overall strength of the QPP during a scan, we used the median of QPP correlation with the scan at supra-threshold local maxima (Fig. 1). Similarly, the median of the time between successive supra-threshold local maxima indicates how often a QPP occurs throughout the functional scan. Hence, these metrics were calculated for 470 low-moderate movers. The frequency characteristics of the QPP correlation timecourse were also examined using the Fourier Transform.

### Comparing QPPs between sessions

The four HCP resting state scans were acquired over two subsequent days with two ~15min back-to-back scans per day, providing an opportunity to examine the relationship between QPPs in the same individual on separate days. QPP analysis was performed separately on the concatenated scans from each day. QPPs were identified based on the modified method introduced earlier, and histograms of correlation with LPCC were calculated for each QPP. As before, the values of the last bin were used as the input to cluster the QPPs of all subjects of both days using Kmeans. QPP type, its median correlation and its periodicity were compared between the two days.

Calculating the correlation between the most representative QPPs of separate days requires both templates to have the same phase; for instance, both start around zero at timepoint 1 and reach their maximum during the first half of their cycle rather than one reaching its minimum in the first half and maximum in the second half. In order to phase-adjust a representative QPP, its correlation time course was cross-correlated with that of all other templates corresponding to all other inspected starting timepoints (see **Fig. S8a).** Templates with values greater than 0.9 were kept since they are similar to the most representative QPP though they can have different phases. These templates were sorted, in descending order, based on their sum of correlation at supra-threshold local maxima, the same metric we have previously used to identify the most representative QPP. The first template whose left early visual area (parcel 184) had the following conditions was designated as the phase-adjusted QPP: 1. near-zero value at timepoint 1,2. average of the first three timepoints is positive, 3. maximum occurring before timepoint 15 and before its minimum, 4. value of the maximum greater than the value of the minimum. If no such template was found, conditions 1 and 2 were discarded and only 3 and 4 were enforced. This method found a phase-adjusted QPP for 465 individuals before GSR, and 467 individuals after GSR. Furthermore, when comparing two QPPs, we performed a fine phase-matching (see **Fig. S8b and S8c),** by time-shifting one of them from −7 to 7 timepoints and taking the maximum Pearson correlation across different time-shifts. For the within subject comparisons, QPPs from each day were compared. For between subject comparisons, QPP of both days from each subject were compared to those of all other subjects, -resulting in 465×464×4 comparisons (467×466×4 after GSR).

### Basic demographics

Out of 470 low-moderate movers, 241 are female (out of 820 HCP with four complete resting state scans, 453 are female). There are 124 twins,62 pair, 31 monozygotic, 196 two-sibling individuals,98 pair, twin or not, 105 three-sibling and 20 four-sibling individuals, two of 3 or 4 siblings might be twin; only 149 individuals have no siblings. The 470 low-moderate movers’ median age is 28 with standard deviation of 3.6 and range of 22-36 (median age of all 820 subjects is 29, standard deviation of 3.7 and range: 22-37).

## Results

Despite the use of the modified method for identifying QPPs, the overall spatial and temporal distribution of activity was similar to that described in Majeed et al., 2011. In all cases, activation and deactivation of areas belonging to the DMN occurred. For some individuals, the TPN exhibited anti-correlation with the DMN, while for other individuals, positively correlated activity was observed in most areas. After global signal regression, however, the QPPS become more similar in that anti-correlation between the DMN and TPN was observed for all individuals. Core regions of DMN include posterior cingulate, precuneus, medial prefrontal, ventral anterior cingulate, lateral parietal, inferior temporal, and parahippocampal areas and core regions of TPN include dorsolateral prefrontal, supramarginal gyrus, posterior parietal, insula, premotor and supplementary motor areas. While all analysis was performed on the individual basis for this manuscript, the resulting spatiotemporal patterns were in good agreement with previous work that calculated QPPs on a group basis (Majeed et al., 2011; Kiviniemi et al. 2016).

**Fig. 2a** shows QPPs from two representative subjects out of the 40 low movers before global signal regression; **Videos 1 and 2** show all 30 timepoints and **Fig. 2b** shows the parcel-wise format. The QPP from the first subject exhibits clear anti-correlation between the activity levels in the DMN and TPN (anti-correlated type of QPP). In the second subject, however, most of the brain activates, with only a few time points showing anti-correlation (most-correlated type of QPP). The histograms of correlations with the LPCC for the two QPPs from the same subjects are plotted in **Fig. 3a.** Within subject 1’s QPP, ~40% of ROIs are anti-correlated with LPCC between −1 and −0.8 while within subject 7’s QPP, no ROI is anti-correlated with LPCC and %75 of the ROIs have a correlation value greater than 0.8. This difference is clearly demonstrated in the correlation maps in **Fig. S9a.** Note, although a few timepoints are showing anti-correlation within Subject 7’s QPP, including all of their 30 timepoint timeseries in calculating the Pearson correlation with LPCC renders a positive but weak correlation. This can be seen by comparing Fig. 2a, timepoint 15 with **Fig. S9a** and is also qualitatively suggested by the parcel-wise format of QPP in Fig. 2b. **Fig. S9b** shows histograms of correlations within the QPPs for all 40 low movers. It is notable that for some individuals, strong anti-correlated activity appears, while for others, most areas are strongly correlated. **Fig. 3b,** in which 40 low movers are sorted based on the percentage of ROIs correlated<-0.8 with the LPCC (left) or sorted based on the percentage of ROIs correlated>0.8 with LPCC (right), summarizes this feature. The clear separation into two groups of data-points in the right part of Fig. 3b provides further evidence that QPPs should be coarsely categorized into two types. Color indicates QPP type and the dashed line indicates where zero is reached. It can be seen that strong anti-correlation exists in some of the individuals categorized as having the most-correlated QPP type in a very small portion of areas. Furthermore, not all individuals categorized as having the anti-correlated type of QPP exhibit broad strong anti-correlations. These findings both point back to the fact that categorization is coarse.

**Figure 2.**
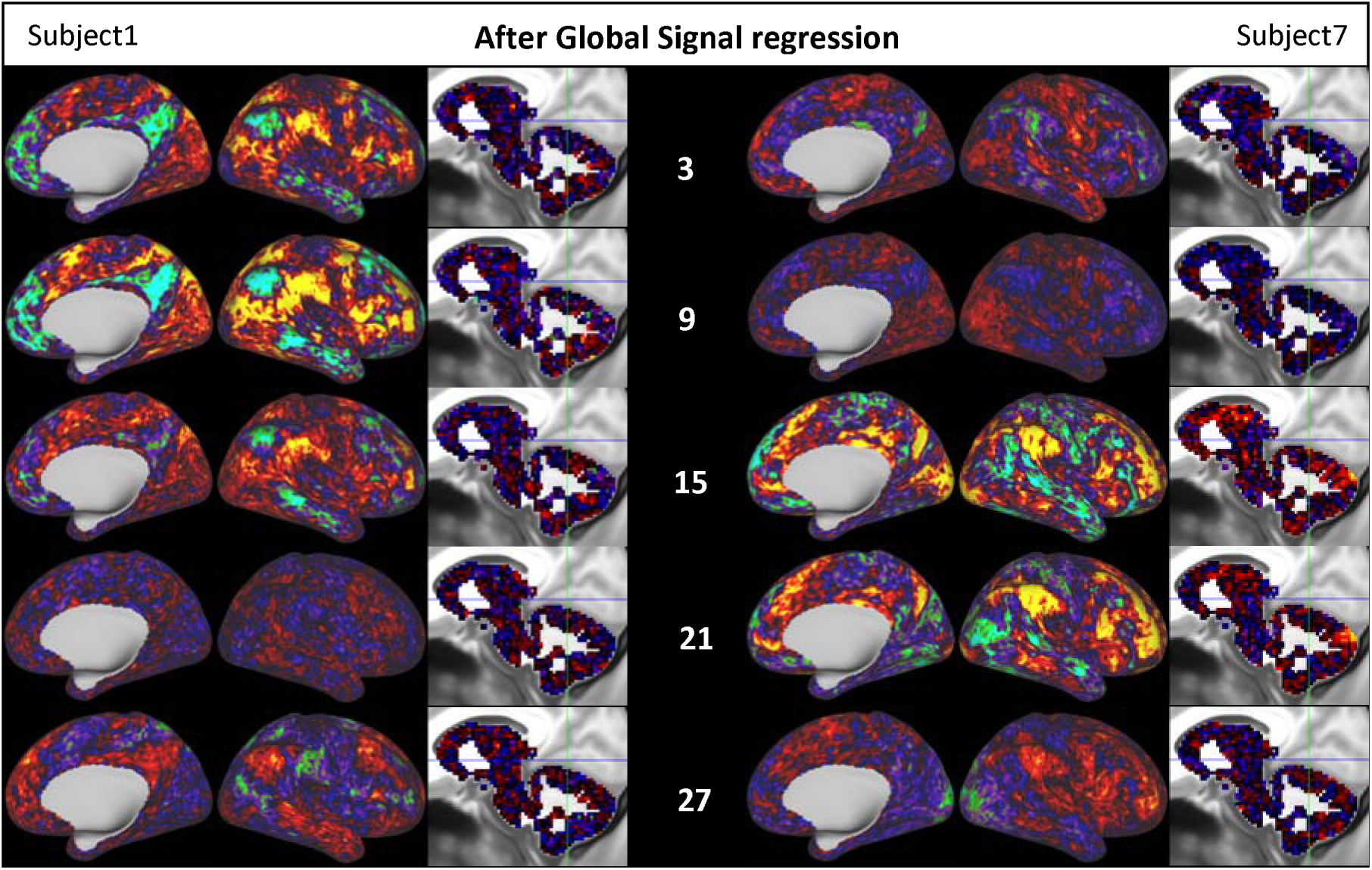
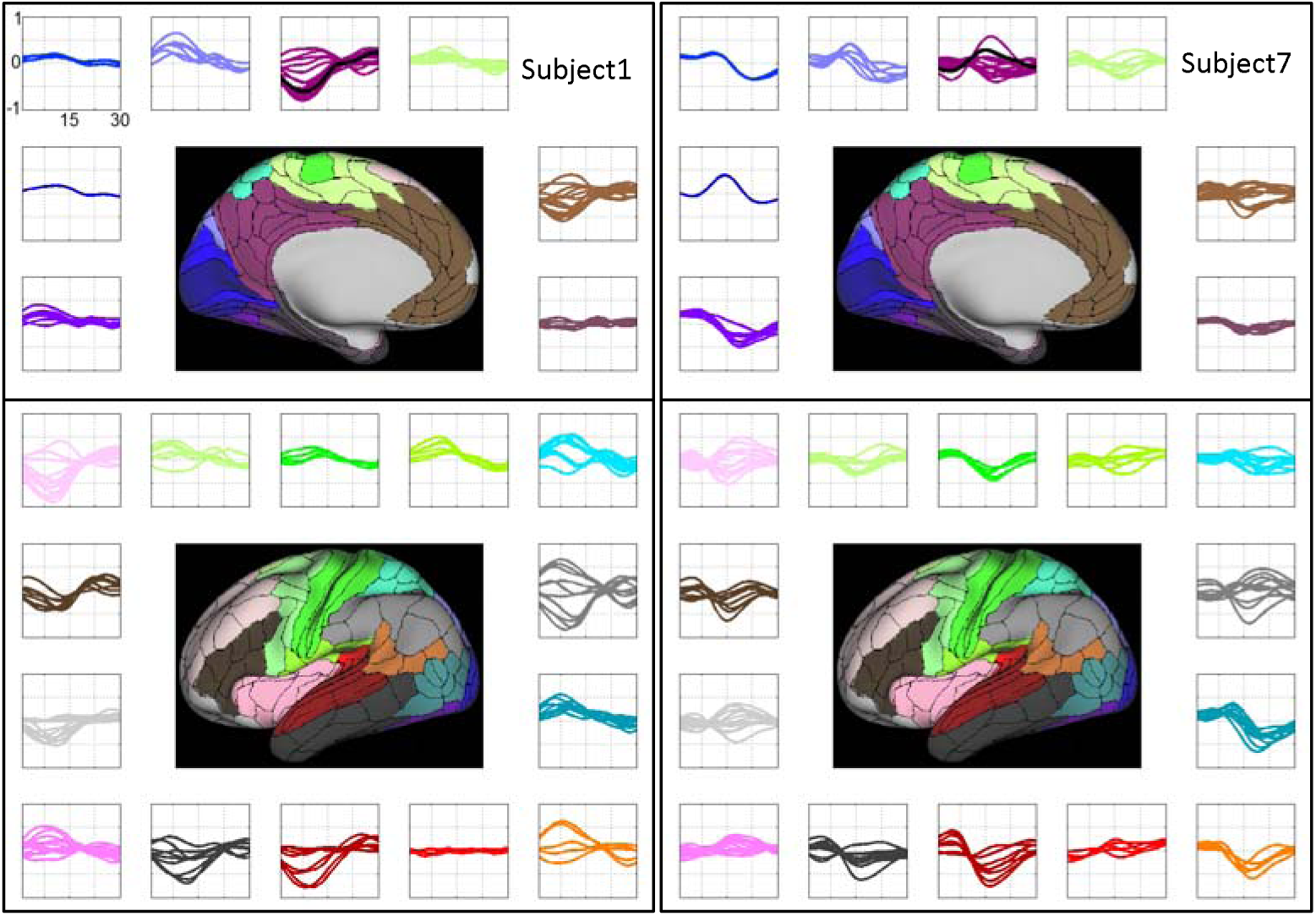
**(a)** QPPs from two representative subjects: One exhibits anti-correlated activity between DMN and поп-DMN areas (anti-correlated type QPP) and the other exhibits correlated activity in most areas, with only a few time points showing anti-correlation (most-correlated type QPP). **(b)** The same subjects shown in parcel-wise format. Subplots showing the signal in the QPP template for each parcel at each time point, with each parcel residing in one of 22 subplots corresponding to Glasser’s 22 Networks (color-coded). **(c)** and **(d)** the same as (a) and (b) after global signal regression. After global signal regression, both subjects exhibit anti-correlation between the DMN and TPN in their templates.

**Figure 3.**
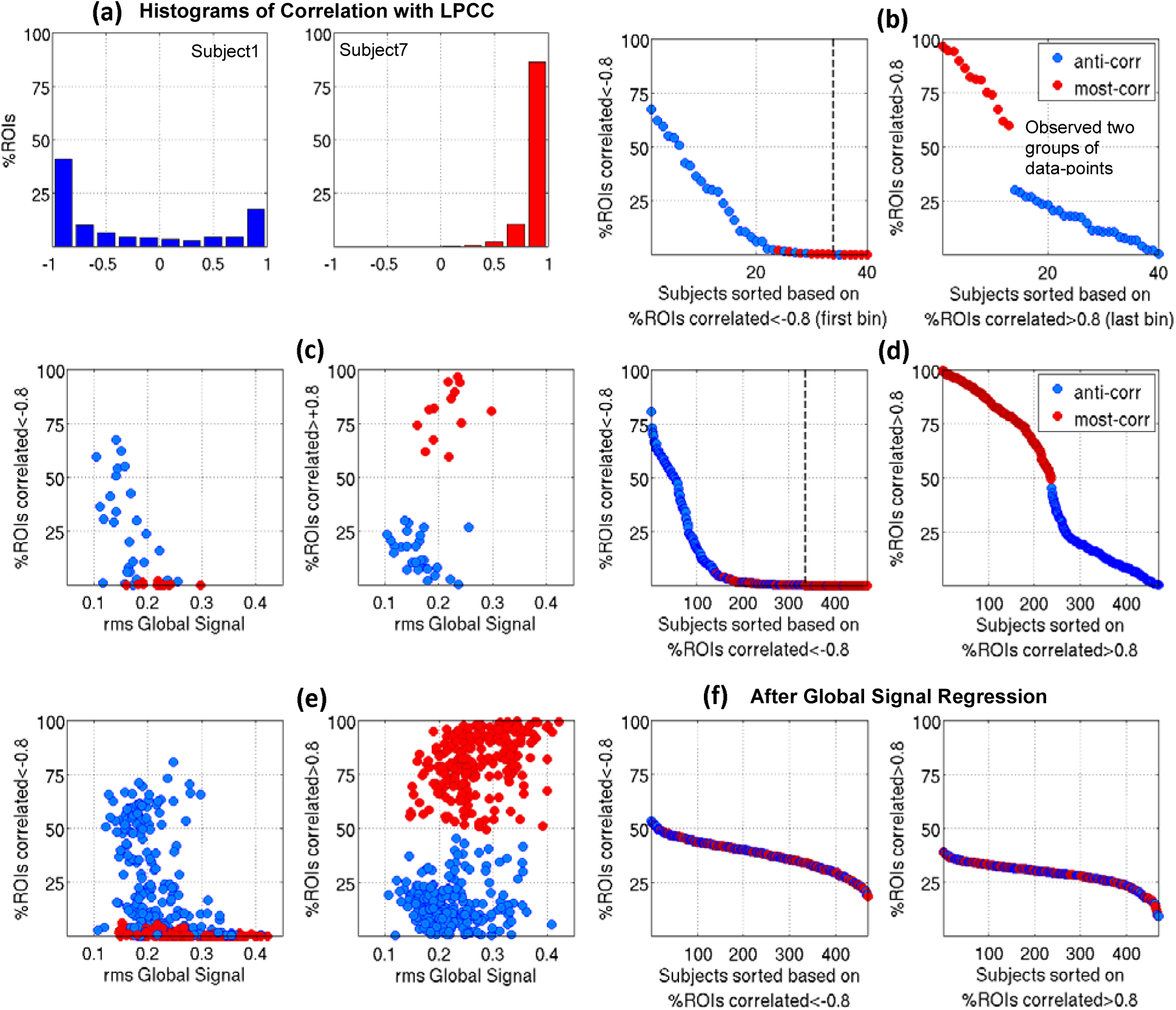
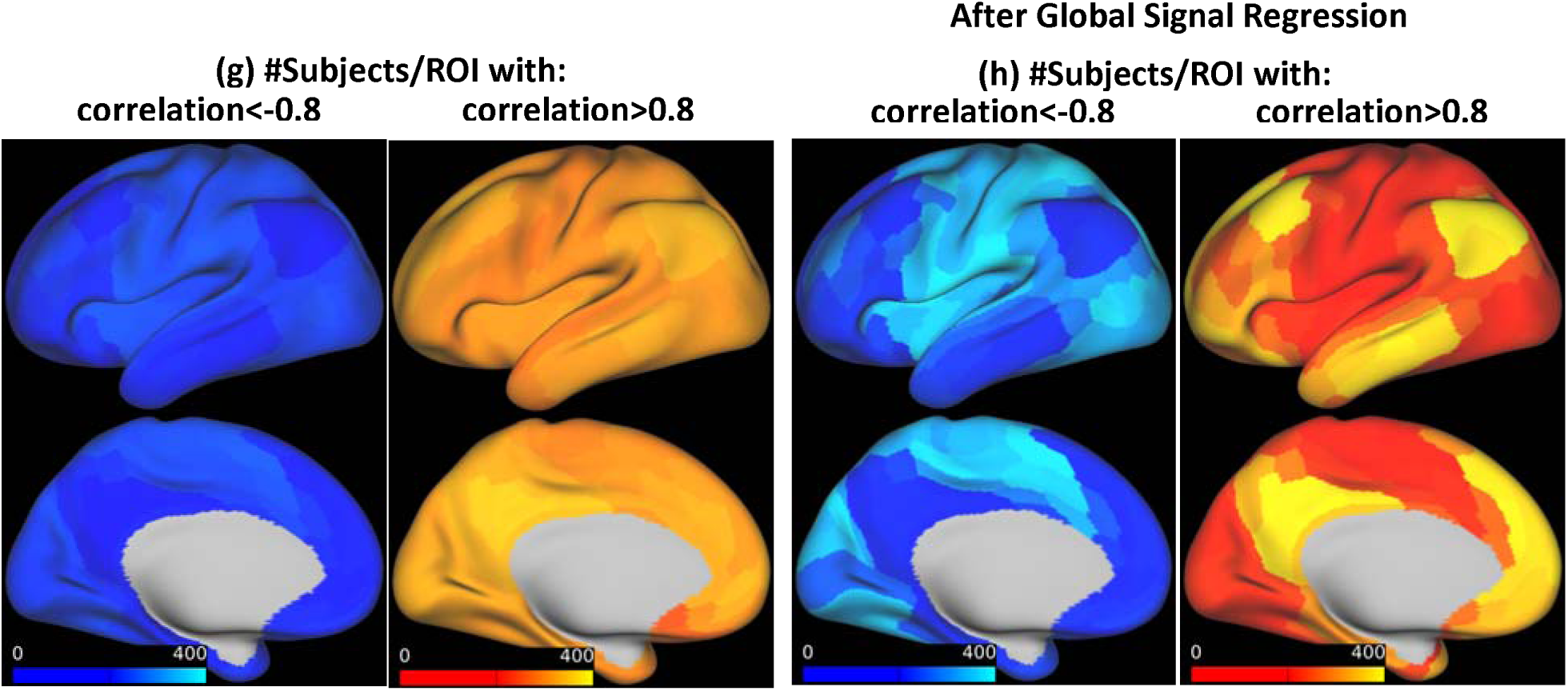
**(a)** histograms of correlations with LPCC for the two QPPs shown in Fig. 2a and b. **(b)** Left plot: 40 low movers sorted based on %ROIs correlated<-0.8 (the value of the first bin of their histogram of correlation), with the dashed line indicating when zero is reached. Right plot: 40 low movers are sorted based on %ROIs correlated>0.8 (the value of the last bin of their histogram of correlation). Two groups of data-points are evident and color indicates QPP type, **(c)** %ROIs with strong negative and positive correlations shown in (b) are plotted against rms of the global signal, **(d) and (e)** are similar to (b) and (c), respectively, for 470 low-moderate movers and show a similar separation into two groups, **(f)** Sorted histograms’ bins similar to (d) but with global signal regression performed before QPP analyses. All 470 subjects show substantial anti-correlation within their QPPs. **(g)** Number of subjects with strong negative and positive correlations per each ROI. Most ROIs with strong anti-correlation belong to non-DMN networks and most ROIs with strong correlation belong to DMN, **(h)** Similar to **(g)** but after global signal regression. Strong correlation and anti-correlation are much more localized.

The widespread involvement of brain areas in the QPPs encouraged us to seek a relationship with the global signal. We plotted the percentage of ROIs with strong negative and positive correlations against the root mean square (rms) of the global signal in **Fig. 3c.** The lower the rms of the global signal, the larger the spatial extent of strong anti-correlations within a QPP, and the higher the rms of global signal, the larger the spatial extent of strong positive correlations within a QPP. The rms of global signal is significantly different between the two QPP types (medians: 0.16 and 0.22, p:2e-4; see whisker plot in **Fig. S9c).**

Similar results, summarized in **Fig. 3d and 3e,** were seen after the pool of subjects was expanded to include individuals with low to moderate levels of motion and analysis was performed on a parcel-wise rather than voxel-wise basis. The rms of the global signal is significantly different between two groups (medians: 0.21 and 0.27, p:7.3e-22, **Fig. S9d).** The number of subjects with strong negative and positive correlations for each ROI are plotted in **Fig. 3g.** As expected, most ROIs with strong anti-correlation belong to поп-DMN networks and most ROIs with strong correlation belong to the DMN. Furthermore, in line with Chen et al., 2017, the seed that maximized the spatial extent of anti-correlations across the majority of subjects is inferior and ventral in LPCC (See **Fig. S10).**

The spatial extent of strong negative and positive correlations within a QPP were plotted versus motion metrics for low movers **(Fig. S11a)** and low-moderate movers **(Fig. S11b and S11c).** Neither of the FD metrics was significantly different between groups with different QPP types; however, the DV metric was slightly although significantly higher in individuals with the most-correlated type of QPP (medians: %8 versus %9, p:3e-4. These findings show that low to moderate levels of motion have minimal impact on the QPP. While higher levels of global signal reduce the amount of anti-correlation observed between the DMN and TPN, the relationship between the global signal and the QPP is not tied to motion but rather to some other aspect of the global signal.

After global signal regression, all 470 subjects showed strong anti-correlations within their QPPs **(Fig. 3f)** in areas belonging to non-DMN networks **(Fig. 3h). Fig. 2c** shows QPPs of the same two subjects in Fig. 2a, but after global signal regression **(Videos 3 and 4** show all 30 timepoints and **Fig. 2d** shows the parcel-wise format). Anti-correlated activity still exists in subject 1 and is now observed in subject 7. Global signal regression makes the detected QPPs more homogeneous across subjects in terms of the spatial extent of strong negative and positive correlation.

To determine whether the occurrence of QPPs coincides with global signal fluctuations in a timepoint-by-timepoint manner, we convolved the QPP of each of the 470 low-moderate movers with its template correlation timecourse. Because the strength of the QPP varies across parcels, for each subject, we considered QPP⊗C at the parcel whose raw timeseries had the highest correlation with the global signal (ROI_gs_). We cross-correlated this timeseries with the global signal (allowed lag: −20 to 20 timepoints). **Fig. S12a** shows the distribution of the results. The median is 0.79 and it is significantly higher than the null, indicating that the QPP does often coincide with the global signal fluctuations in a timepoint-by-timepoint manner with small time-shifts. Moreover, the higher the rms of the global signal, the stronger this coincidence. **Fig. S12b** shows the number of subjects exhibiting the highest correlation with the global signal for each parcel; as expected, primary and early visual, somato-motor and auditory areas are the most correlated with the global signal in agreement with (Glasser et al., 2016) and (Power et al., 2017). In addition to taking QPP⊗C at ROI_GS_, three alternatives were tested: (i) selecting parcel 184, Left Early Visual Area, for all subjects, since this was the parcel with the highest correlation with the global signal overall **(Fig. S12d),** (ii) selecting the parcel at which cross-correlation of QPP⊗C and the global signal was maximum **(Fig. S12f),** and (iii) averaging QPP⊗C across parcels and cross-correlating that with the global signal **(Fig. S12e).** Results were similar for all three approaches.

**Fig. 4a** shows the rms of the global signal plotted against the standard deviation of respiratory variation (RV) and heart rate variation (HV). Fluctuations in the global signal are positively correlated with respiratory and heart rate variations, in line with Power et al., 2017 where at low motion timepoints, global signal and RV correlation was shown to be ~0.5. RV and HV are themselves correlated **(Fig. S13a).** The spatial extent of strong negative or positive correlations within a QPP is solidly predicted by rms of the global signal; hence, they should also be related to respiratory variations. As expected, individuals with the anti-correlated QPP type have slightly, but significantly, lower RV and HV variation compared to those with most-correlated QPP type **(Fig. S13b).**

To further examine the relation between QPPs and slow physiological variations, estimates of slow respiratory- and cardiac-induced BOLD signal fluctuations were regressed from the data and the residuals were reanalyzed. **Fig. 4b** shows that more subjects now exhibit the anti-correlated QPP type. Out of 292 subjects whose physiological noise estimates were regressed, 156 had the anti-correlated type of QPP, and after regression this number increased to 225. Moreover, the spatial extent of strong anti-correlations has increased overall (compare **Fig. 4b** with **Fig. 3d)** in areas that belong to non-DMN networks **(Fig. S14).** The beta value obtained from regression, the cross-correlation values and their corresponding lags that preceded regression stage to determine the optimal lag per parcel are provided in **Fig. S15.** Overall, in our dataset, the BOLD signal correlated more with the respiration variation, in more widespread areas, compared to cardiac variations. Primary areas of visual, auditory, and somato-motor cortex had the highest correlation with respiratory and cardiac variation, but the correlation in default mode areas was not prominent and was less than in neighboring areas.

**Figure 4.**
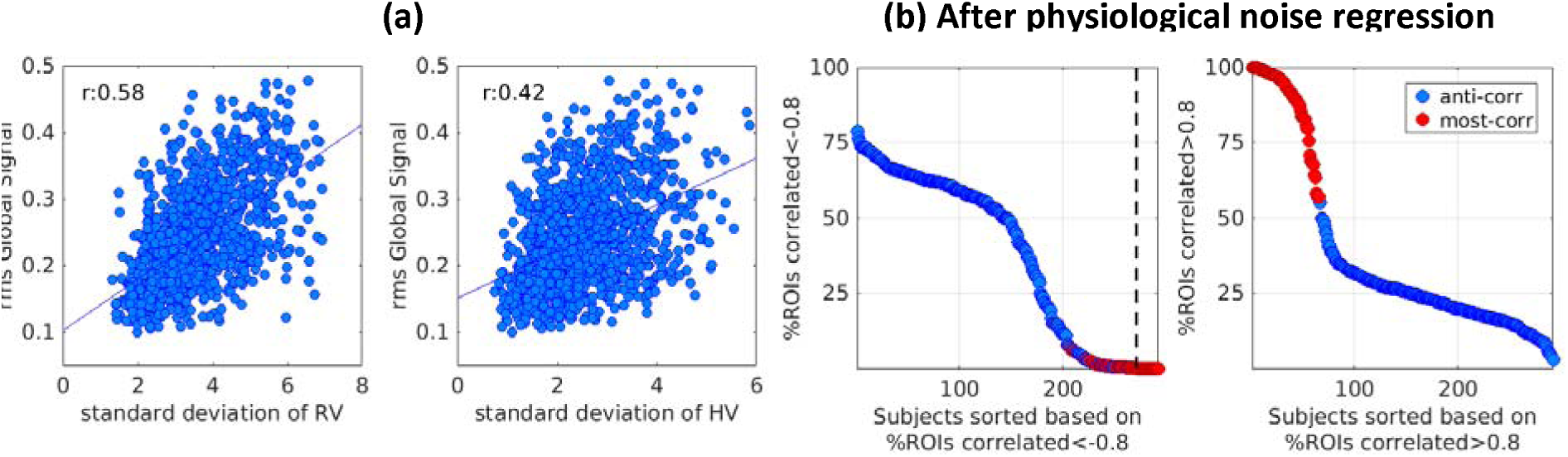
**(a)** rms of global signal is correlated with standard deviation of respiratory variation and heart rate variation (4×292 datapoints correspond to four scans of 292 individuals included). **(b)** Subjects sorted based on the first and the last bin of the histogram of correlation within their QPPs, after physiological noise regression; more subjects exhibit anti-correlated type QPP.

How correlated is the QPP with the functional scan and how often does it occur? The answer can be obtained by inspecting the correlation time course for the QPP and the scan. **Fig. 5a** shows such time courses for the two representative subjects in Fig. 2. Subject1’s QPP (anti-correlated type), correlates less with subject1’s functional scan as opposed to subject7’s QPP (most-correlated type). Also, subject1’s QPP occurs less often, with more variability relative to that of Subject7’s. For all 470 subjects, the median of QPP correlation at the supra-threshold maxima was 0.35 and the median of the time between successive maxima was 26s **(Fig. 5b).**

**Figure 5.**
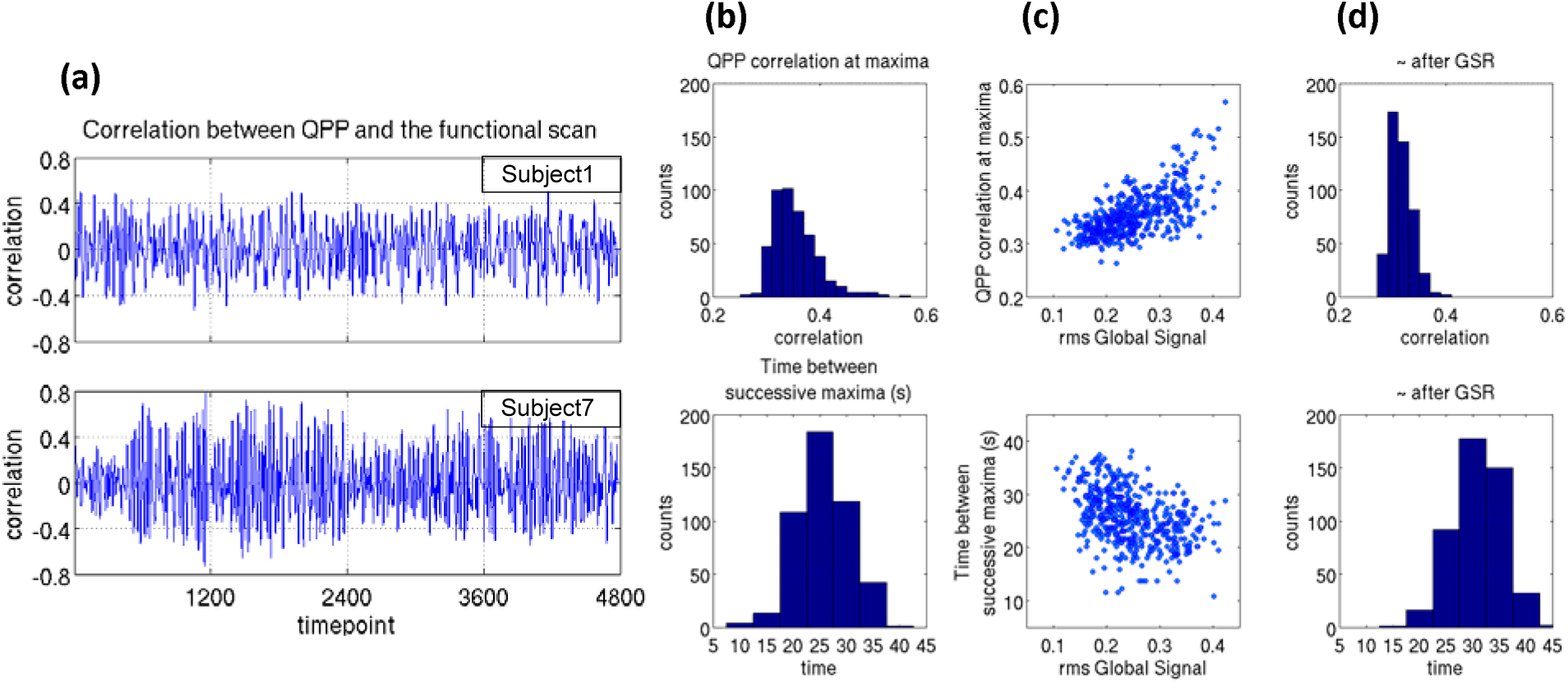
**(a)** The time course of correlation between QPP and the functional scan for the same two representative subjects in Fig 2a and 3a. The median of QPP correlation at supra-threshold local maxima, which describes the strength of the QPP, is 0.31 for Subject 1 and 0.36 for Subject7. The median of the time between successive maxima, which describes how often the pattern occurs, is 30s and 22s, respectively (the sum of all QPP correlation values at supra-threshold local maxima, which depends on both the strength of the pattern and how often it occurs, is 32 for Subject 1 and 57 for Subject 7). As observed at the group level, in these two representative subjects, the QPP occurs more often and is more strongly correlated in the subject with higher global signal (Subject 7). **(b)** For all 470 subjects, histograms describe the median of QPP correlation at supra-threshold local maxima (top, median:0.35, range:0.26-0.57) and time between successive supra-threshold local maxima (bottom, median:26s, range:11-38s). **(c)** values of (b) versus rms of global signal; a higher rms of the global signal (likely the most-correlated QPP type) corresponds to higher correlation and periodicity, **(d)** same as (c) after global signal regression; QPPs’ correlation and periodicity decrease (correlation has median of 0.31 and range of 0.27-0.4, and time difference has median of 32s and range of 15-44s).

**Fig. 5c** shows values of Fig. 5b versus rms of global signal, the predictor of strong negative or positive correlations within a QPP. High rms of the global signal corresponds to high correlation and periodicity, meaning most-correlated type QPP correlates relatively more with the functional scan and occurs more often. Low rms of global signal corresponds to low correlation and periodicity, meaning, anti-correlated type QPP correlates relatively less with functional scan and occurs relatively less often. After global signal regression, where all QPPs have strong anti-correlation within them, correlations and periodicities decrease **(Fig. 5d,** medians:0.31 and 32s). This behavior was expected since after GSR all QPPs become similar to the anti-correlated QPP type before GSR.

To demonstrate the quasi-periodic nature of the QPPs, we plotted the average Fourier transform of all individuals’ QPP correlation time courses in the frequency domain as shown in **Fig. 6.** A broad peak at ~0.03Hz (~30s) provides evidence of the quasiperiodic nature of the pattern.

**Figure 6.**
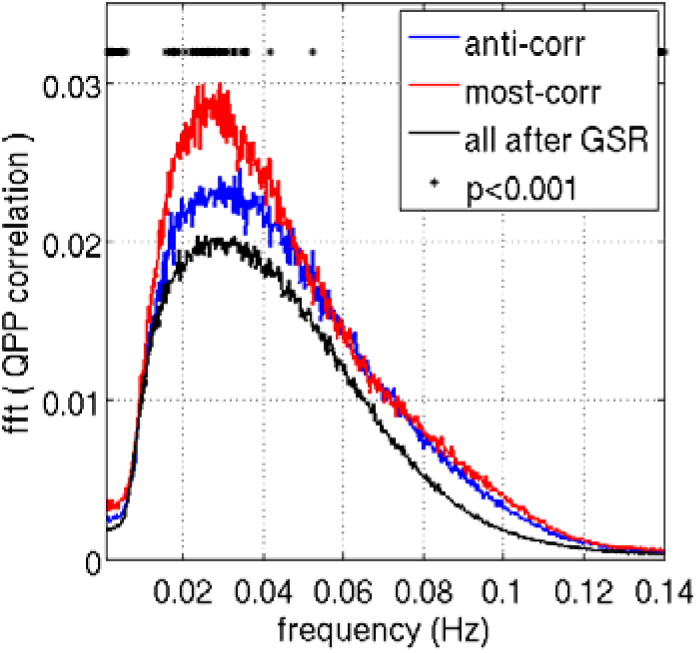
The averaged Fourier transform of QPP correlation across all subjects exhibits a broad peak at 0.02-0.04 Hz, qualitative evidence of the quasi-periodic nature of the pattern. The peak is significantly different between the two QPP types or when comparing before and after global signal regression.

To examine the relationship between QPPs of the same individual on subsequent days, QPP analysis was performed separately on the concatenated scans of each day. Out of the 245 individuals with the anti-correlated type QPP on day 1, 178 (~%73) exhibit the same type on day 2, and out of the 225 individuals with most-correlated type QPP on day 1, 130 (~%58) exhibit the same type on day 2 **(Fig. 7a);** this suggests individuals with anti-correlated type QPP are more stable in their QPP type. The rms of the global signal, which predicts QPP type, is highly correlated between days (r=0.79, **Fig. 7b),** suggesting a very interesting point that global signal fluctuation can be regarded as a trait, in addition to reflecting brain states (Wong et al., 2013). Also note that the standard deviation of Respiration and Heart rate Variation which is correlated with global signal (r=0.58 and 0.42, Fig. 4a) is reasonably correlated between days (r=0.56 and 0.58, **Fig. S16a),** which is in line with Birn et al., 2014. QPP correlation and periodicity are collectively reflected in the sum of QPP correlation at supra-threshold local maxima which is reasonably correlated between days (r=0.61, **Fig. 7c).** Both the median of QPP correlation at supra-threshold maxima and the median of the time between successive maxima remain correlated between days (r=0.43 and 0.44, **Fig. S16b).** QPP correlation and periodicity are still correlated across days after global signal regression **(Fig. S16c)** although the correlation values decrease, suggesting that these metrics could potentially be recognized as traits independent of the global signal. **Fig. 7d** shows that phase-adjusted QPPs, derived based on the procedure explained in the methods section, are significantly more correlated between days within subjects (mediamo.78) than between subjects (median of 0.65; p_KStest_: 1.4e-88). This finding is unchanged by global signal regression, although medians are reduced **(Fig. S16d,** within subject: 0.63, between subject:0.42, p_KStest_:1.4e-88).

**Figure 7.**
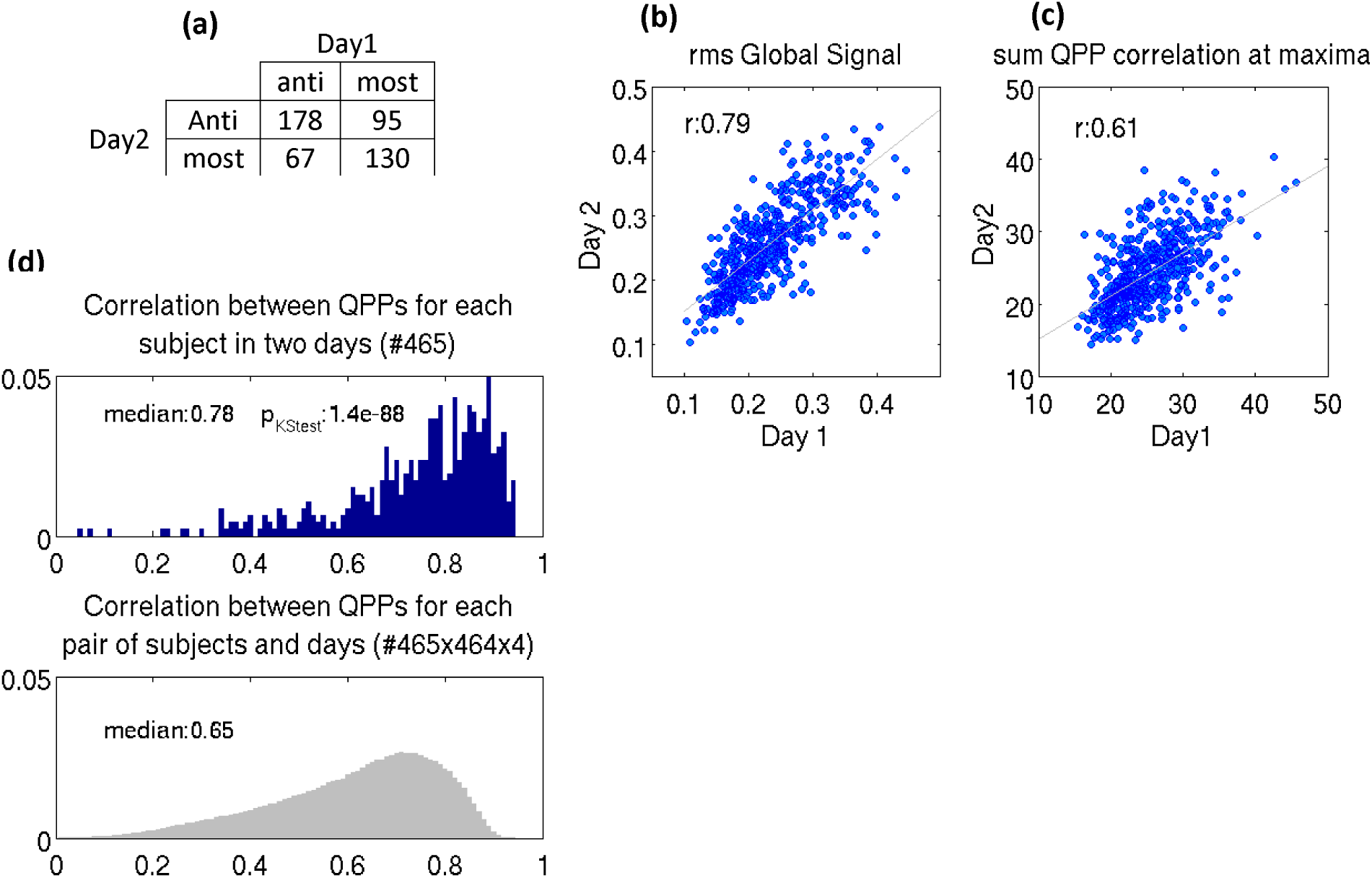
Between two days: **(a)** individuals with anti-correlated type QPP are more stable in their QPP type, **(b)** the rms of the global signal is highly correlated, as is **(c)** QPP correlation and periodicity collectively reflected in the sum of QPP correlation at supra-threshold maxima, **(d)** QPPs are significantly more correlated across days within subjects than between subjects (area under distributions has been normalized to 1).

Out of 233 individuals with anti-correlated type QPP, 148 are female (~%63.5) and out of those with most-correlated type QPP, %60.8 are male. Out of 31 monozygotic twins, 20 have the same QPP type (%64.5); this percentage is %61.3 for 31 non-monozygotic twins. Out of 196 two-siblings individuals (twin or not) %46 have the same QPP type; this percentage is %29 for 105 three-sibling individuals and %20 for 20 four-sibling individuals (two of 3 or 4 might be twins). There is no age difference between group of individuals with two different QPP types (both medians are 28, p=0.85).

## Discussion

### Summary

In order to perform the first examination of the variability in QPPs across individuals, we developed a robust version of the pattern-finding method that does not depend upon a randomly-chosen starting point. Applying this method to ~500 individuals, we found that QPPs fall into two coarse categories prior to global signal correction and that these categories are closely linked to the overall level of global signal. After global signal regression, QPPs become remarkably similar in their spatial extent of strong positive or negative correlation, strength and timing across individuals. The QPPs do not appear to result from motion or physiological signal fluctuations, suggesting a neuronal source. Between subsequent days, QPP type, strength and periodicity are reasonably correlated and the pattern itself is somewhat individual-specific.

### Potential significance of QPPs

The involvement of the DMN and TPN suggests that QPPs may be linked to attention or vigilance. The DMN is defined by (i) deactivation during tasks requiring attention to the external environment, and (ii) activation during internally focused and self-referential tasks (Buckner et al., 2008). The TPN is defined as areas that co-activate during tasks requiring externally focused attention. It follows that the DMN and TPN tend to be anti-correlated (Fox et al., 2005; Murphy and Fox 2016). One can further speculate, based on the definitions of the two networks, that the activity levels in each network prior to a task could influence subsequent activity when a task commences, hence predicting measures of behavioral-level performance. Numerous works support this hypothesis (see Rosenberg et al., 2016; Tavor et al., 2016; Smith et al., 2015; Cole et al., 2014; Schultz and Cole 2016; Sadaghiani et al. 2015, 2010, 2009). In addition to predicting cognitive traits, a preliminary QPP analysis during a psychomotor vigilance task shows that the phase of the QPP predicts reaction time and can therefore reflect varying cognitive states even within an individual (Abbas et al., 2016b).

Moreover, because numerous neurological and psychiatric disorders exhibit alterations in the DMN in particular (see Broyd et al., 2008 for a comprehensive review), it is plausible that some of these changes may be reflected in alterations in the QPP. By revealing the course of activation and deactivation of DMN, QPP has the potential to add to our basic cognitive science or improve diagnosis and treatment assessment in mental disorders and neuro-pathological diseases.

### QPPs and physiological rhythms

Physiological noise is known to contribute spatially-structured noise into functional connectivity measurements and could conceivably contribute to the large-scale, repetitive patterns observed in the QPPs. Slow variations in respiration depth and rate give rise to fluctuations in the arterial level of C0_2_, a potent vasodilator, causing variations in cerebral blood flow and oxygenation hence variations in the BOLD signal (Birn et al., 2006; Chang et al., 2009). Heart beat (rate and contractility) is tightly coupled with respiration (rate and depth) in both fast and slow varying regimes (Power et al., 2017; Chang et al., 2009). However, our findings suggest physiological variations impact the type of QPP, but they do not account for its presence, which makes it more likely that the QPP is of neuronal origin. Furthermore, Kiviniemi et al., 2016 utilized the QPP algorithm to examine processes explicitly associated with cardiac pulsation and respiration in Magnetic Resonance Encephalography (MREG) data with very high temporal resolution (100ms), and the spatiotemporal patterns that they obtained are much different in spatial extent and timing than the QPPs reported here.

In Power et al. 2017, Respiration Volume per unit Time (RVT), introduced by Birn et al., 2006, explained more variation of the global signal compared to RV, so we calculated RVT as well (procedure described in **Fig. S17a).** RV and RVT are correlated **(Fig. S17b)** and both capture respiration variation in rate and depth. The rms of global signal is correlated with standard deviation of RVT (r:0.47, **Fig. S17c),** but the correlation is lower than for RV (r:0.58), hence, we did not pursue further analyses based on RVT.

### Global Signal

Global signal regression is a controversial topic (Power et al., 2017; Murphy and Fox 2016; Caballero-Gaudes and Reynolds 2017). Sustained motion-induced and respiratory-induced signal fluctuations are largely and simply removed by global signal regression, but it may also remove possible widespread neuronal co-activations, increase the spatial extent and strength of anti-correlations, and distort the variation structure (Saad et al., 2012). In this work, we have solely quantified individual differences in the fluctuation level of the global signal, partially explained by differences in the fluctuation levels of physiological rhythms, and observed that global signal regression leads to more homogenous spatial extent of strong negative and positive correlations within the QPPs of individuals. Note that the QPP algorithm is a correlation-based approach and using it on the whole brain is likely to identify the pattern with the strongest activity and largest spatial extent. Without GSR, the QPP algorithm is biased toward global fluctuations which are strong and extensive in those individuals who exhibit such fluctuations and unsurprisingly, their QPPs turn out to be the most-correlated type. After GSR, the next largely extended activation pattern throughout the whole brain is the anti-correlated activity of DMN and TPN and therefore the QPPs turn out to be the anti-correlated type.

A feature of the global signal is worth highlighting: its distribution is bimodal **(Fig. S18a)** and the borderline between two modes approximately matches the borderlines in Fig. 3e that separate mostly correlated QPPs from QPPs with strong anti-correlations, providing additional evidence that QPPs, which identify the global patterns, should be coarsely categorized into two types.

Lastly note the rms values of global signal reported here are based on the average of the timeseries with our three additional preprocessing steps applied. These rms values are highly correlated (r:0.95) with rms values without any additional preprocessing steps named MGT (mean of grayordinate timeseries) in the HCP dataset **(Fig. S18b).**

### Motion

Motion has complex effects on the MRI signal that are only partially compensated with correction strategies. While it seemed unlikely that the repetitive patterns detected with the QPP algorithm could arise from head motion, we used a subject group with very low motion in our original analysis. The results were similar to previous studies of QPPs and suggest that motion is not a major contributor to the QPP. After this original analysis, the subject pool was expanded to include a much larger number of subjects with more typical levels of motion. The 470 subjects of this study were chosen based on an arbitrary threshold on FD **(Fig. S2a)** and neither of two FD metrics were significantly higher in individuals with the most-correlated QPP type relative to those with the anti-correlated type **(Fig. S11).** Moreover, we performed a supplemental analysis of the 40 highest movers and found that the results were very similar to the low-moderate movers **(Fig. S19).** Based on these findings, motion does not appear to be a major issue for our future work on QPPs using the HCP dataset. Nevertheless, motion should be considered when evaluating differences in QPP metrics across groups.

### Limitation and Future work

The QPP algorithm was initially developed to capture repeated large scale patterns of brain activity. Work in rodents that limits the algorithm to small sections of the brain (e.g., the subcortical grey matter) suggests that other, less prominent patterns may coexist with the QPP described in this manuscript (Majeed et al., 2011), an avenue that should be explored in future work. The current work focuses on characterizing the most representative QPP and how it varies across individuals. Subcortical regions were included only in the group of 40 low movers where analysis was performed on a voxel-wise basis because the parcellation scheme that we used for larger groups is limited to the cortex. However, based on the voxel-wise analysis (Fig. 2), subcortical areas exhibit potentially interesting patterns of activation and deactivation during the QPP, and we intend to investigate their involvement further during future work.

To achieve practical computation times while using the modified QPP method that examines every time point, analysis was performed on a parcel-wise basis for most of this work. Our preliminary analysis, based on 40 low movers, has shown that QPP type, correlation histogram and relation with the global signal, do not change when analyzed using cortical parcels as opposed to cortical and subcortical voxels. Also, as shown in Glasser et al., 2016b or Chen et al., 2015, parcel-wise analysis has the additional advantage of increasing the statistical significance. In our results, for instance, QPP correlation significantly increases. However, some of the fine details of the QPP are lost due to the spatial downsampling. For example, the propagation along the cortex observed by Majeed et al. 2009 and 2011 or even in this report (particularly evident in the videos) is difficult to detect in the parcel-wise QPP.

For the current analysis, all of the free parameters for the QPP analysis were kept constant across subjects. However, the outcomes suggest that some of these parameters should be optimized on an individual basis. For example, visual inspection of all 470 QPP1s shows that the full cycles are slightly different from 30 timepoints for some individuals (see **Fig. S20** for a typical and two extreme examples). As shown in Majeed et al., 2011, the QPP algorithm is robust in finding templates whose duration slightly deviates from a preset value, and qualitatively all the results presented in this work, such as QPP type and relation with global signal, seem robust in this regard as well, but systematically determining the optimal window-length for each individual is a goal of our future work.

The modified QPP algorithm uses all time points to avoid any potential biases. Nevertheless, it is clear from Fig. 1b that considerable redundancy results. A carefully-selected subset would ensure robust detection of the primary QPP while reducing the computation time. The selection of this subset will be explored in future work.

The origins of the QPPs remain poorly understood and are critical to understanding the neuro-physiological processes that are represented in MRI-based measurements of functional connectivity (FC). The spatially coherent fluctuations of the QPP certainly contribute to FC measurements, while the relatively long time scale and repetitive nature of the patterns is evidence that they are not solely serving cognitive processing attributed to intrinsic activity of DMN and TPN. To determine if QPPs were linked to the variations in correlation observed with sliding window techniques, we looked at phase amplitude coupling between infra-slow activity and higher frequencies (Thompson et al, 2014b), but found little evidence of a relationship. Further work showed that infra-slow electrical activity correlated with QPPs and higher frequency activity correlated more closely to sliding window BOLD correlation (Thompson et al, 2015), strong evidence that the QPPs and remaining BOLD signal have differing sensitivities to neural activity and do not reflect the same underlying processes. The link between QPPs and infra-slow electrical activity suggests that QPPs are likely to reflect a distinct brain process and at a minimum, they are a source of neuro-physiological BOLD fluctuations that can be removed as a nuisance signal like respiration or cardiac pulsation. Both the residual BOLD FC maps and the QPP templates that characterize intrinsic activations of DMN and TPN have the potential to provide new insight into the basic organization of the brain as well as neurological and psychiatric disorders. Examining the behavioral correlates of various features of the QPP is an especially crucial avenue for future work that will help us understand the role it plays in the brain’s functional organization.

## Conclusion

This work provides the first assessment of the variability in individual QPPs and their relationship to global signal and physiological parameters. Improving the characterization of QPPs might provide further insight into the organization of dynamic brain activity, which in turn could underlie behavioral differences in the healthy population or connectivity changes in neurological and psychiatric disorders.

## Acknowledgment

The authors thank Derek Smith, Anzar Abbas, and Dr. Savannah Cookson for their comments and Dr. Shiyang Chen and Dr. Jacob Billings for their very helpful discussions. We also thank Dr. Matt Bezdek, Dr. Wenju Pan, Dr. Garth Thompson, Amrit Kashyap, Prof. David Van Essen, Dr. Cesar Caballero-Gaudes, Dr. Bruce Crosson, and Alican Nalci for their one-time yet valuable comments. This work was funded by the NIH grants R01MH1 11416-01 and R01NS078095 and NSF grant BCS INSPIRE 1533260.

